# The autoactivity of tomato helper NLR immune proteins of the NRC clade is unaltered in *prf* mutants of *Nicotiana benthamiana*

**DOI:** 10.1101/2025.03.11.642614

**Authors:** Daniel Lüdke, Hsuan Pai, AmirAli Toghani, Adeline Harant, Chih-Hang Wu, Sophien Kamoun

## Abstract

Nucleotide-binding domain and leucine-rich repeat immune receptors (NLRs) can function in networks of sensors and helpers to induce hypersensitive cell death and immunity against pathogens. The tomato sensor NLR Prf guards the Pto kinase from AvrPto and AvrPtoB effector perturbation and activates the downstream helpers NRC2 and NRC3. Prf is conserved across the Solanaceae and its ortholog in the model species *Nicotiana benthamiana* is also required for detection of AvrPto/AvrPtoB function on Pto. A recent study reported that cell death induction after transient expression of an autoactive mutant of tomato NRC3 is abolished upon RNAi silencing of *Prf* in *N. benthamiana*. Here we generated loss-of-function *prf* mutants in *N. benthamiana* and demonstrate that autoactive mutants of eight canonical tomato NRCs (NRC0, NRC1, NRC2, NRC3, NRC4a, NRC4b, NRC6, and NRC7) still induce hypersensitive cell death when expressed transiently in the *prf* mutant background. Autoactive tomato NRCs also triggered cell death when expressed in lettuce (*Lactuca sativa*), an Asteraceae plant that does not have a *Prf* ortholog. These results confirm a unidirectional dependency of sensors and helpers in the NRC network and underscore the value of the *N. benthamiana* and lettuce model systems for studying functional relationships between paired and networked NLRs.

## Introduction

Intracellular recognition of pathogen effector proteins by NLRs typically leads to induction of cell death (Jones and Dangl, 2006). While singleton NLRs can detect effectors and induce cell death, paired and networked NLRs can be distinguished into phylogenetic and functional clades of sensor and helper NLRs (Contreras et al., 2023a). In the NRC (NLR required for cell death) network of asterid plants, sensors detect effectors and induce the oligomerization of downstream helpers. Activated helpers form membrane localized resistosomes, which act as calcium channels to induce cell death (Wu et al., 2017; Ahn et al., 2023; Contreras et al., 2023b; Liu et al., 2024; Madhuprakash et al., 2024). The activation of NRC helpers by sensors follows the activation-and-release model, in which sensors are not part of activated helper resistosomes (Ahn et al., 2023; Contreras et al., 2023b; Madhuprakash et al., 2024). The cell death function of helpers is mediated by the N-terminal coiled-coil domain, encoding the conserved MADA motif (Adachi et al., 2019), while sensors have degenerated N-termini and can encode an N-terminal solanaceous-domain (SD), which is present in a subclade of NRC sensors (Contreras et al., 2023a).

NRC helpers are widely present across Solanaceae and can be grouped into 11 distinct phylogenetic sub-clades (Lüdke et al., 2023; Madhuprakash et al., 2024). Several disease resistance genes are NRC sensors which require NRCs from the helper clades for cell death induction and immunity. Previously studied NRC network components include the sensors Rpi-amr1a and Gpa2 which signal through NRC2/3, Rpi-amr1e, Rpi-amr3, Bs2, Rx, R1, Sw-5b, and R8 signaling through NRC2/3/4, as well as Rpi-blb2 and Mi-1.2 which exclusively signal through NRC4 (Wu et al., 2017; Witek et al., 2021; Lin et al., 2022; Lin et al., 2023). While the Hero resistance gene signals specifically through NRC6 (Lüdke et al., 2023), the pepper (*Capsicum annum*) *Ca*Rpi-blb2 signals through NRC8/9 (Oh et al., 2023). The *Pseudomonas syringae* pv. tomato (*Ps*t) resistance protein Prf from tomato (*Solanum lycopersicum*) contains an SD-domain, which is required for interaction with the Pto kinase (Mucyn et al., 2006; Ntoukakis et al., 2014). The Prf/Pto complex binds and detects the *Ps*t effectors AvrPto and AvrPtoB, leading to activation of NRC2 and NRC3 for cell death induction and immunity (Wu et al., 2004; Wu et al., 2016; Wu et al., 2017; Wu and Kamoun, 2021; Sheikh et al., 2023; Zhang et al., 2024). A Prf ortholog is present in the model species *N. benthamiana* and is also required for detection of AvrPto/AvrPtoB function on Pto (Lu et al., 2003; Mucyn et al., 2006).

A recent study reported that cell death induction after transient expression of *Sl*NRC3^H478AD479V^, an autoactive mutant of *Sl*NRC3, is abolished upon RNAi silencing of *Prf* in *N. benthamiana* (Zhang et al., 2024). This work suggests that the cell death of autoactive NRC3 is genetically dependent on the Prf sensor, in sharp contrast with the previously reported unidirectional network architecture (Wu et al., 2017). Here, we revisited these experiments using mutant *N. benthamiana* plants rather than RNAi. We used CRISPR/Cas9 to generate three independent loss-of-function *prf* mutant lines to determine the extent to which autoactive mutants of helper NLRs of the NRC phylogenetic clade can induce cell death independently of *Prf*. In addition, we expressed autoactive helper NRCs in lettuce (*Lactuca sativa* var. Fenston), an Asteraceae plant that does not encode for a *Prf* homolog (Pai et al., 2025). Our results revealed that autoactive variants of eight tomato NRC helpers, including *Sl*NRC3, can still trigger cell death in the absence of functional Prf. These findings confirm the unidirectional nature of the NRC sensor-helper networks of NLRs and demonstrate the utility of *N. benthamiana* genetic mutants and lettuce as a model system for dissecting sensor-helper interactions.

## Results and Discussion

The *N. benthamiana* genome encodes two *Prf* copies, *Prfa* (Niben101Scf00650) and *Prfb* (Niben101Scf01984) (Kourelis et al., 2021; Supplementary Fig. S1). We used CRISPR/Cas9 to generate *N. benthamiana* lines with loss-of-function mutations in both copies, targeting the first 300 nucleotides, respectively (Fig. 1A). These loci encode a predicted coiled-coil within the SD-domain (Supplementary Fig. S2), important for Pto interaction (Saur et al., 2015). While *prf* 1-4 contains an in-frame deletion in *Prfa* and *Prfb, prf* 1-5 and *prf* 2-1 contain frame shifts in *Prfa* and *Prfb*, respectively (Fig. 1A). To test the *prf* lines for AvrPto recognition, we performed Agrobacterium-mediated transient expression. While AvrPto, and Pto/AvrPto expression leads to cell death in wild type plants, no cell death was observed in *prf* mutant lines (Fig. 1B, Supplementary Fig. S3). Loss of cell death was complemented by co-expressing *Sl*Prf with Pto/AvrPto, while neither Pto, nor *Sl*Prf expression induced cell death (Fig. 1B). Co-expression of Rx with *Potato virus X* coat protein (CP), was used as cell death control induced by an NRC sensor-helper pair. These results indicate that all generated mutant lines, including the in-frame *prf* 1-4 mutant, are *Prf* loss-of-function mutants.

**Figure 1.**
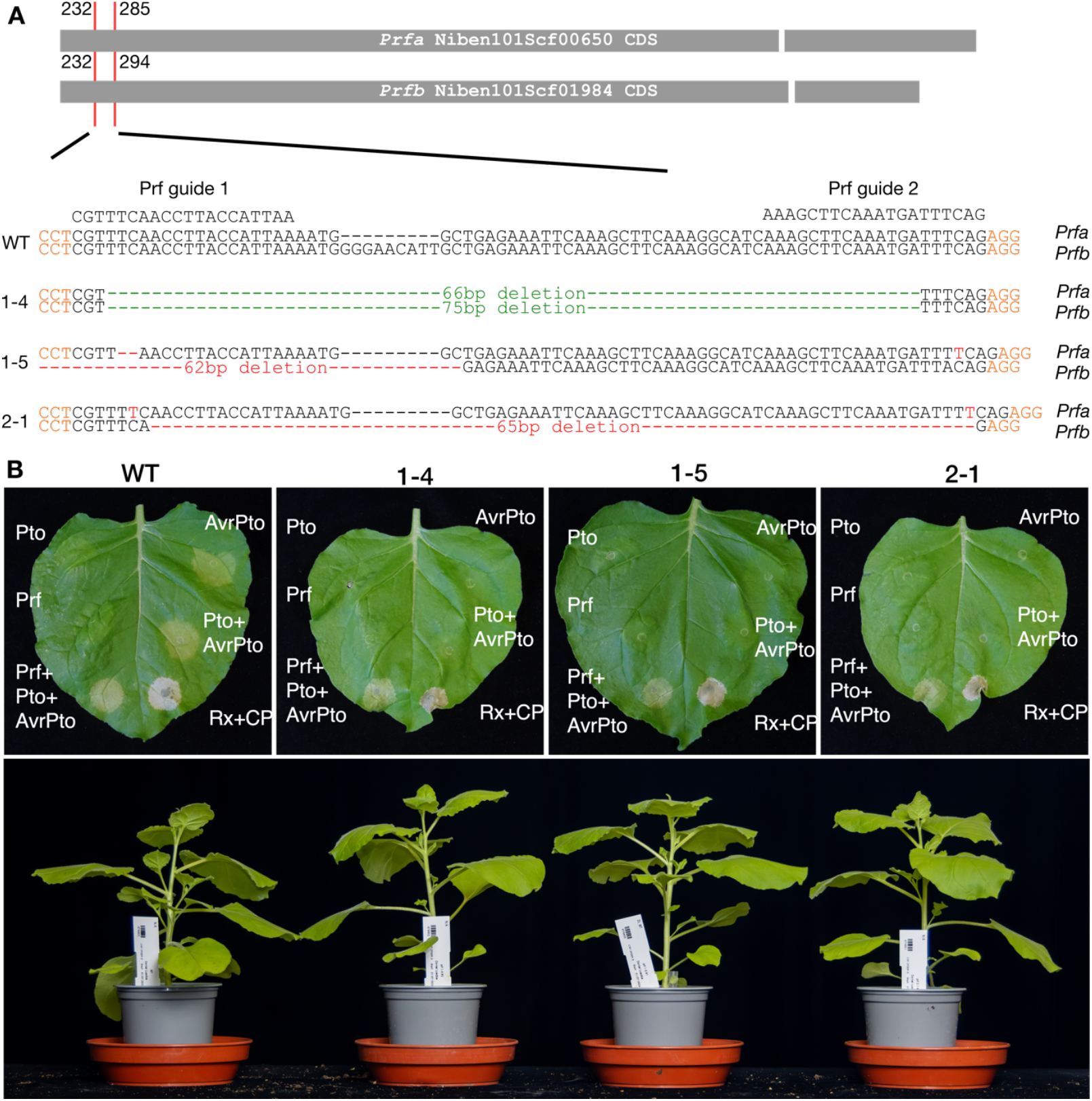
CRISPR/Cas9 generated *prf* mutant lines do not respond to AvrPto or Pto/AvrPto. **(A)** Schematic overview of *Prfa* and *Prfb* CDS targeted by CRISPR/Cas9. The CDS of *Prfa* and *Prfb* from *N. benthamiana* is indicated as gray bars, drawn to scale. Breaks between bars indicate the position of introns. The red lines indicate the position of guideRNA targets in the CDS. The numbers indicate the start and end nucleotide position targeted by the guideRNAs within the CDS. The wildtype (WT), *prf* 1-4, *prf* 1-5, and *prf* 2-1 nucleotide sequence within the target region of *Prfa* and *Prfb*, received by amplicon sequencing, is shown. Orange indicates protospacer adjacent (PAM) motifs, red indicates frame shift nucleotide deletions (−) or insertions, green indicates deletions in-frame. **(B)** Representative cell death phenotypes in *N. benthamiana* wildtype (WT), *prf* 1-4, *prf* 1-5, and *prf* 2-1 leaves induced by transient expression of AvrPto, Pto/AvrPto, Prf/Pto/AvrPto, or Rx/CP (top). Complete quantification and statistical analyses are presented in Supplementary Fig. S3. Representative image of wildtype (WT), *prf* 1-4, *prf* 1-5, and *prf* 2-1 growth phenotypes of 6-week-old plants grown in a glasshouse (bottom).

The tomato reference genome encodes a total of 11 proteins in the NRC phylogenetic helper clade (NRCX, NRC0, NRC1, NRC2, NRC3, NRC4a, NRC4b, NRC4c, NRC5, NRC6, and NRC7), all of which have canonical signatures of functional helper NLR proteins, except for NRC5, which does not encode an MHD motif (Lüdke et al., 2023) and NRCX, which acts as a modulator protein (Adachi et al., 2019). We tested if transient expression of the 9 canonical tomato NRC helpers as autoactive MHV mutant variants leads to the induction of a cell death response in the *prf* mutant lines. Expression of tomato NRC0, NRC1, NRC2, NRC3, NRC4a, NRC4b, NRC6, and NRC7 as autoactive MHV mutant variants induced a cell death response in the *prf* mutant lines similar to the wildtype plants (Fig. 2A). In contrast to results received by RNAi silencing of *Prf* in *N. benthamiana*, we could not detect a significant reduction of cell death after transient expression of autoactive versions of NRC helpers in *prf* mutant lines (Fig. 2A, Supplementary Fig. S4). Next, we tested if autoactive tomato NRC mutant variants induce a cell death response when expressed transiently in lettuce (*Lactuca sativa*), which does not encode a *Prf* homolog (Pai et al., 2025). Similar to transient expression in *N. benthamiana*, tomato NRC0, NRC1, NRC2, NRC3, NRC4a, NRC4b, NRC6, and NRC7 induced a cell death response in lettuce when expressed as autoactive MHV mutant variants, but not as wildtype protein (Figure 2B). We conclude that autoactive NRC helper proteins can function in the absence of functional sensor NLRs like Prf, consistent with the activation-and-release model (Ahn et al., 2023; Contreras et al., 2023b).

**Figure 2.**
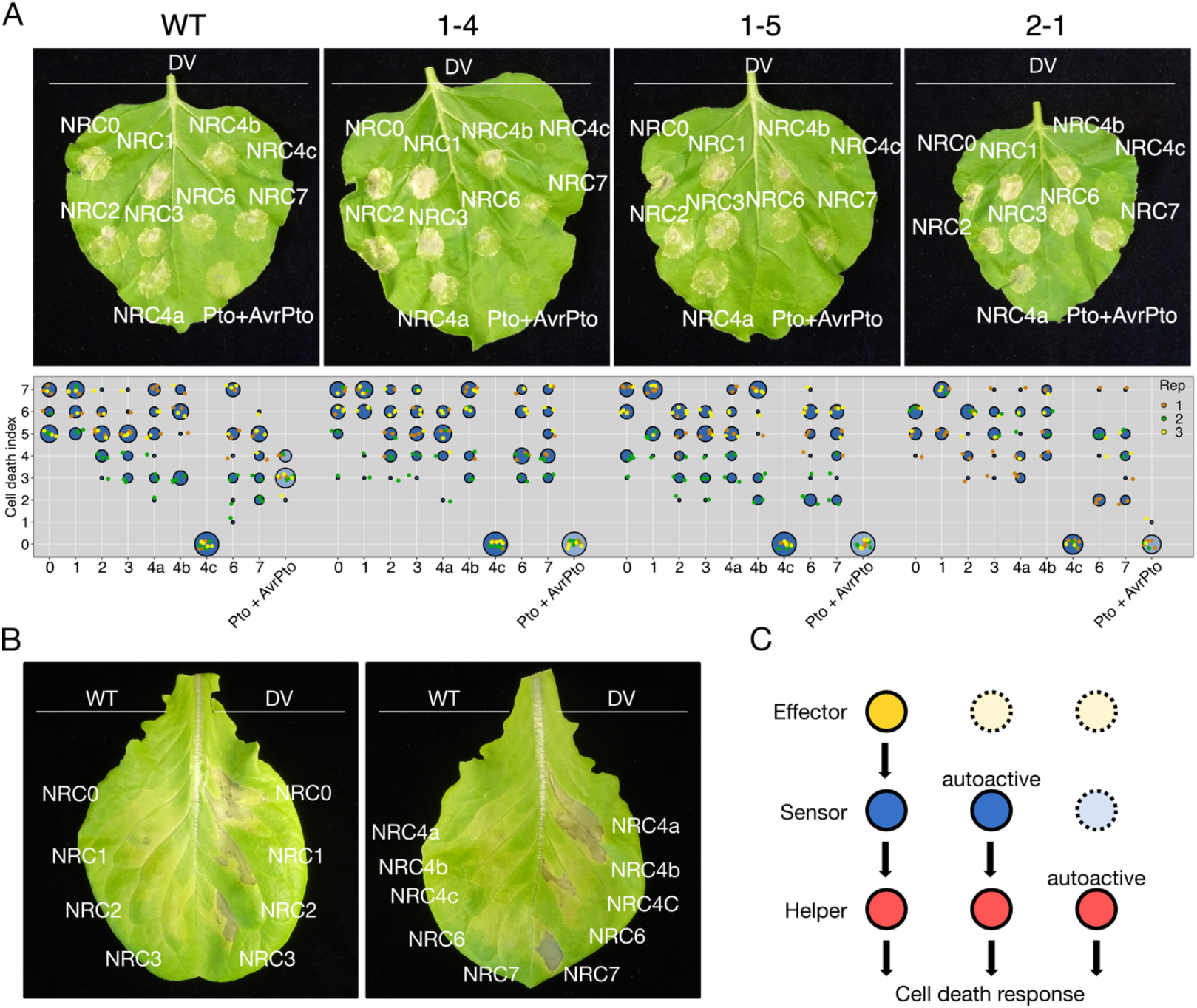
Cell death induced by autoactive tomato NRCs is unaltered in *N. benthamiana prf* mutants and lettuce. **(A)** Representative cell death phenotypes in *N. benthamiana* wildtype (WT), *prf* 1-4, *prf* 1-5, and *prf* 2-1 leaves induced by transient expression of autoactive MHV mutants of tomato NRCs, photographed at 7 days post-infiltration with agrobacteria. Cell death data quantification from the leaves is displayed below the leaves. The cell death index for each individual spot on the leaves is represented as dots, with different colors for biological replicates. The central circle for each cell death category proportionally indicates the total number of data points. Statistical analysis is shown in Supplementary Fig. S3. **(B)** Representative cell death phenotypes in leaves of *L. sativa* var. Fenston induced by transient expression of autoactive MHV mutants of tomato NRCs, photographed at 7 days post-infiltration with agrobacteria. **(C)** Schematic representation of directionality for cell death signaling in the NRC network. Dotted lines indicate that the component is no longer required for the induction of a cell death response. While effector activated or autoactive sensors require a downstream helper NRC for the induction of cell death, autoactive helper NRCs can act independently of upstream sensors or effectors for the induction of a cell death response.

Several studies outlined that Prf signals through NRC2 and NRC3 for the induction of cell death and immunity. We showed that NRCs can induce cell death when expressed as autoactive mutants in the absence of functional Prf, either in *N. benthamiana* or in lettuce. Our results are consistent with the activation-and-release model (Ahn et al., 2023; Contreras et al., 2023b) and the observation that activated Prf sensors and NRC2/NRC3 helpers do not form a stable complex (Sheikh et al., 2023). In addition, NRC-dependent sensors are not part of NRC helper hexameric resistosomes (Contreras et al., 2023b; Madhuprakash et al., 2024). Our phylogenomic analyses are also consistent with the view that NRCs do not require Prf to function. Whereas *NRC3* is present in all 35 examined Solanaceae genomes, *Prf* genes are missing in six of the 35 Solanaceae genomes, further suggesting that *Prf* is dispensable for NRC3 function (Supplementary Fig. S1). In addition, autoactive tomato NRC mutants can trigger cell death when expressed in lettuce, which does not carry a Prf or an NRC3 ortholog and diverged from tomato 97.5 – 109.2 MYA (Pai et al., 2025).

Altogether, these findings are consistent with the view that the edges in NRC network of sensors and helpers have unidirectional relationships. While effector activated or autoactive sensors require a downstream helper for the induction of cell death, autoactive helpers can act independently of upstream sensors for the induction of a cell death response (Figure 4c).

## Acknowledgments

We thank Suomeng Dong (Nanjing Agricultural University) for valuable discussions and for carefully reading the manuscript. We are grateful to Mark Youles and Liam Egan from the Sainsbury Laboratory Synbio support team for generating the CRISPR/Cas9 vector used. We thank Matthew Smoker, Jodie Taylor, and Aleksandra Wawryk-Khamdavong from the Sainsbury Laboratory Tissue Culture & Transformation support team for transformation of *Nicotiana benthamiana*.

## Author contributions

Conceptualization: D.L., S.K.

Data Curation: D.L., A.T., H.P.

Formal Analysis: D.L., A.T., H.P.

Investigation: D.L., A.T., H.P., A.H.

Methodology: D.L., A.T., H.P., A.H.

Resources: D.L., A.T., H.P., A.H., C.H.W.

Software: D.L., A.T.

Supervision: D.L., S.K.

Funding Acquisition: S.K.

Project Administration: D.L., S.K.

Writing Initial Draft: D.L.

Editing: D.L., A.T., S.K.

## Supplementary Data

Supplementary Materials

Supplementary Figure S1: Phylogeny of the Prf clade

Supplementary Figure S2: AlphaFold3 prediction of NbPrfa and NbPrfb

Supplementary Figure S3: Cell death quantification and statistics for Figure 1B

Supplementary Figure S4: Cell death quantification and statistics for Figure 2A

Supplementary File 1: Primers used in this study

Supplementary File 2: Plasmids used in this study

Supplementary Dataset 1: Phylogenetic analyses of the Prf clade

Supplementary Dataset 2: CRISPR/Cas9 vector maps

Supplementary Dataset 3: Amplicon sequencing raw reads

Supplementary Dataset 4: Cell death quantification images and data

Supplementary Dataset 5: AlphaFold3 generated PDB files of NbPrfa and NbPrfb

## Data availability

All data is available in the Supplementary Data of this manuscript or under https://doi.org/10.5061/dryad.sxksn03d6, https://doi.org/10.5281/zenodo.14720919 (Toghani et al., 2025) and https://doi.org/10.5281/zenodo.15000078.

