## Supplementary Data for "The autoactivity of tomato helper NLR immune proteins of the NRC clade is unaltered in *prf* mutants of *Nicotiana benthamiana*"

**Supplementary Data**
**Supplementary Materials**
**Plasmid construction**
The Golden Gate Modular Cloning (MoClo; Weber et al., 2011) and MoClo plant parts kits (Engler et al., 2014) were used for cloning. Wildtype and MHV-mutant variants of tomato NRC helpers were synthesized as *N. benthamiana* codon-optimized L0 modules in pICH41155 through GENEWIZ/AZENTA (https://www.azenta.com/). The Prf (AAF76312.1) CDS was amplified from 35S:Prf-3HA (Mucyn et al., 2006) and domesticated to remove BsaI sites. All gene were either cloned into the binary vector pJK001c (Paulus et al., 2020), pJK268c (Kourelis et al., 2020), or pICH86988 (Weber et al., 2011). All Plasmids used and constructed in this study are described in Supplementary Data 7. To generate guid RNA modules encoding each target site within the genomic *Prf* sequence, Phusion polymerase standard PCR reactions were performed using sgRNA_Prf_1 or sgRNA_Prf_2 as forward and universal_rev as reverse primers (Supplementary File 1) with pICSL70001 as a PCR template. The resulting PCR products were cloned into pICH47751-P3F and pICH47761-P4F, together with pICSL90002 (U6-26) as a promoter fragment, using a GoldenGate cloning reaction (Engler et al., 2014). The sequence verified plasmids were used to assemble the CRISPR vector DL355 (pL2V-NPTII/FCY-UPP/SlFAST/pRPS5a::mEGFP-iCas9-tTriple/dummyP1/dummyP2/sgPrf1/sgPrf2/end-linker) using pICSL22022 (pL2V-NPTII(pICSL11024)/FCY-UPP(pICSL11243)/SlFAST(pICSL11242)/pRPS5a::mEGFP-iCas9(pICSL11244)-tTriple/), pICH54011 (dummyP1), pICH54022 (dummyP2), pICH47751-P3F (sgPrf1), pICH47761-P4F (sgPrf2), and pICH41780 (end-linker) in a GoldenGate cloning reaction. All maps of plasmid templates and of the DL355 plasmid are available as Supplementary Dataset 2.

**Plant growth conditions**
All wildtype and *prf* mutant *N. benthamiana* plants as well as *L. sativa* var. Fenston were grown in Levingtons F2 compost in a glasshouse setting with natural light for cell death assays or genotyping performed.

**Plant transformation**
*N. benthamiana* plants were transformed using *A. tumefaciens* as described (Clemente, 2006). Briefly, 4-week-old *N. benthamiana* leaves of up to 10 cm in diameter were surface sterilized by dipping into 70% ethanol and immersing into 1% sodium hypochlorite solution, containing Tween 20 as surfactant, for 25 minutes. The *A. tumefaciens* pMP90 strain carrying the binary plasmid DL355 was gown, harvested, and washed in inoculation buffer [per liter: 4.3 g MS basal salts (1×), 30 g sucrose]. 1-2 $cm^2$ leaf sections were prepared after rinsing with water and incubated with the *A. tumefaciens* suspension. Leaf sections fully immersed into the agrobacterium inoculation suspension were blotted dry, and transferred to co-cultivation medium [per liter: 4.3 g MS basal salts (1×), 30 g sucrose, 1× Gamborg's B5 vitamins, 0.1 mg Naphthalene Acetic Acid (NAA), 0.59 g MES, pH = 5.7 using KOH, 4 g Agargel] supplemented with 1 mg/L Benzylaminopurine (BAP). Plates were incubated in a plant growth chamber at 24 °C and 18-h light for 3 days, before transferring explants to plates containing shooting media (co-cultivation medium media supplemented with 1 mg/L Benzylaminopurine (BAP), 500 µg/ml Cefotaxime/Augmentin, and 100 µg/ml kanamycin). Explants were subcultured onto fresh shooting media every 7-10 days until the first shoots appeared. Shoots >3 mm were excised and transferred to rooting media [per liter: 2.15 g MS basal salts (0.5×), 5 g sucrose, pH = 5.8 with KOH, 2.5 g Gelrite] supplemented with 500 µg/ml Cefotaxime/Augmentin and 100 µg/ml kanamycin. Explants were

cultured until roots appeared, and T0 plantlets were transferred to soil and grown in the glasshouse as described.

**Plant genotyping**

T0 lines were grown in a glasshouse as described to set seeds. Seeds of the T1 generation were grown on sand drenched with 1/4 MS solution containing 2mM 5-fluorocytosine (5-FC) for negative selection of the FCY-UPP marker (Davis et al., 2009). Germinated Cas9 free plants were transferred to a greenhouse, grown as described and DNA was isolated using the DNeasy Plant Mini Kit (QIAGEN) according to manufacturer's instructions. Phusion polymerase was used in a standard PCR reaction to amplify the CRISPR/Cas9 target region using primers Prf_fwd and Prf_rev (Supplementary File 1). PCR fragments were sent for amplicon sequencing using the GENEWIZ Amplicon-EZ service to determine mutations. Processing and alignment of the received reads was performed using Geneious Prime 2024.10.30 (Kearse et al., 2012). Fastq files containing raw reads were imported as paired reads (inward pointing) and trimmed using BBDuK 38.84 to remove adapter sequences, low quality reads (<20), and reads shorter than 20bp. Merging of trimmed paired reads was performed using BBMerge (Bushnell et al., 2017). Merged reads were aligned to the *Prf* amplicon sequences of *NbPrfa* and *NbPrfb*. Homozygous *prf* mutant plants were used for subsequent experiments performed. Amplicon sequencing raw reads are available as Supplementary Dataset 3.

**Transient gene expression and cell death assays**

Transient gene expression was performed by infiltrating *A. tumefaciens* strain GV3101 pMP90 in *N. benthamiana* or strain C58C1 in lettuce, each transformed with respective binary expression constructs. Strains were inoculated from glycerol stocks and grown O/N at 28°C in LB supplemented with appropriate antibiotics. Cells were harvested by centrifugation at 2000 × g for 10 min at RT, resuspended in infiltration buffer (10 mM MgCl2, 10 mM MES-KOH pH 5.6, 200 µM acetosyringone), and incubated for 2 hours at room temperature. Infiltration into 5 to 6-week-old *N. benthamiana* leaves was performed with a needless syringe using an OD600 of 0.3 for all strains used. Cell death phenotypes for *N. benthamiana* were scored with a range from 0 (no visible necrosis) to 7 (fully confluent necrosis) according to Adachi et al., 2019. Quantification and statistical analysis was performed by using the besthr R library (MacLean, 2019) and plotted using a script described in (Bentham et al., 2023). Scoring for all experiments can be found in Supplementary Dataset 4.

**Phylogenomic analyses**

18,021 NLR sequences were extracted using NLRtracker (Kourelis et al., 2021) on 40 Asterales proteomes. Proteins with the domain architectures "CNL", "CCNL", "CCCNL", "CNLO", "CN", "OCNL", "CONL", "NL", "NLO", "ONL", "BCNL", "BNL", "BCN", "BCCNL", "BNLO", "BOCNL", "BCNLO", "BBCNL", "BBNL", "RNL", "TN", "TNL", "TNLO", and "TNLJ" were retained (15,344 sequences). Sequences with NB-ARC domains exceeding 400 amino acids, or subsiding 250 amino acids were excluded, resulting in a final dataset of 13,179 sequences. NB-ARC domains of the dataset and the reference RefPlantNLR dataset (Kourelis et al., 2021) were aligned using FAMSA v2.2.2 (Deorowicz et al., 2016) and an initial phylogenetic tree was constructed using FastTree v2.1.11 (Price et al., 2010). The NRC superclade, including RefPlantNLR reference NRC helper and sensor sequences, was extracted from a well-supported branch of this tree. A sub tree was generated using FastTree after realigning with MAFFT v7.526 (Katoh and Standley, 2013). Well-supported branches of the sub tree were used to identify and extract the Prf superclade and the NRC helper clade. These sequences were realigned, and phylogenetic trees were reconstructed using MAFFT and FastTree. Subsequently, the Prf and NRC3 clades were extracted from their respective trees, aligned, and trees were generated using MAFFT and FastTree. The Prf and NRC3 clades were used to extract the copy numbers and frequencies of Prf and NRC3 gene homologs across species. All sequence processing steps were conducted in R using the Biostrings package (H. Pagès, P. Aboyoun, R. Gentleman, and S. DebRoy, 2017). Phylogenetic trees were handled using Dendroscope v3.8.1 and visualized using iTOL (Huson and Scornavacca, 2012; Letunic and Bork, 2021). All generated data is available as Supplementary Dataset 1 or under https://doi.org/10.5061/dryad.sxksn03d6 or https://doi.org/10.5281/zenodo.14720919 (Toghani et al., 2025). All scripts are available at https://github.com/amiralito/Prf_NRC3.

**Structural modelling**

AlphaFold 3 [Seeds = 1] was used to generate models of full-length monomeric NbPrfa and NbPrfb (Abramson et al., 2024). The top predicted structure for each protein was used and ChimeraX was used color domains, the corresponding CRISPR deletion regions, and to display pLDDT scores (Pettersen et al., 2021). PDB files of the structural models are available as Supplementary Dataset 5.

**Supplementary Figures**

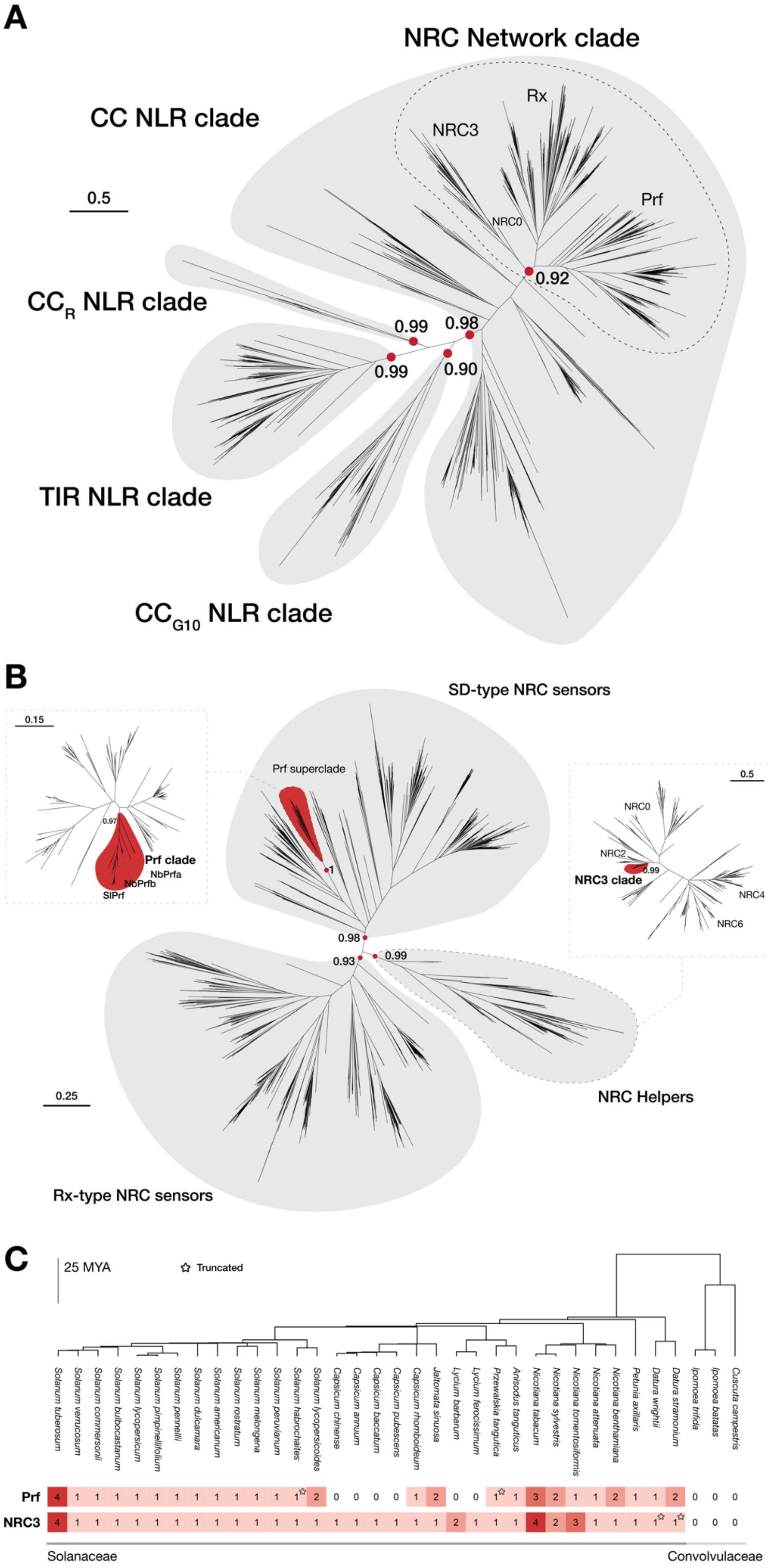

**Supplementary Figure 1: Phylogeny of the Prf clade. (A)** The NB-ARC domain based phylogenetic tree of NLRs was constructed using the NLR sequence database (Toghani et al., 2025). NLR sequences were aligned using FAMSA v2.2.2 and FastTree v2.1.11 was used to determine the phylogeny of the NRC network clade. **(B)** Subtrees for NRC helper and SD-type NRC sensor subclades were generated based on well-defined bootstrap values using MAFFT v7.526 and FastTree. The Prf clade and the NRC3 clade form a distinct branch of the SD-type NRC sensor subclade or the NRC helper subclade, respectively. **(C)** The phylogenetic tree of Solanaceae was constructed using TimeTree, with Convolvulaceae as outgroup. The number of Prf and NRC3 homologs encoded for each species is indicated at the bottom. Truncated sequences are indicated by a star. The divergence time estimated in MYA is based on TimeTree. All data is available as Supplementary Dataset 1.

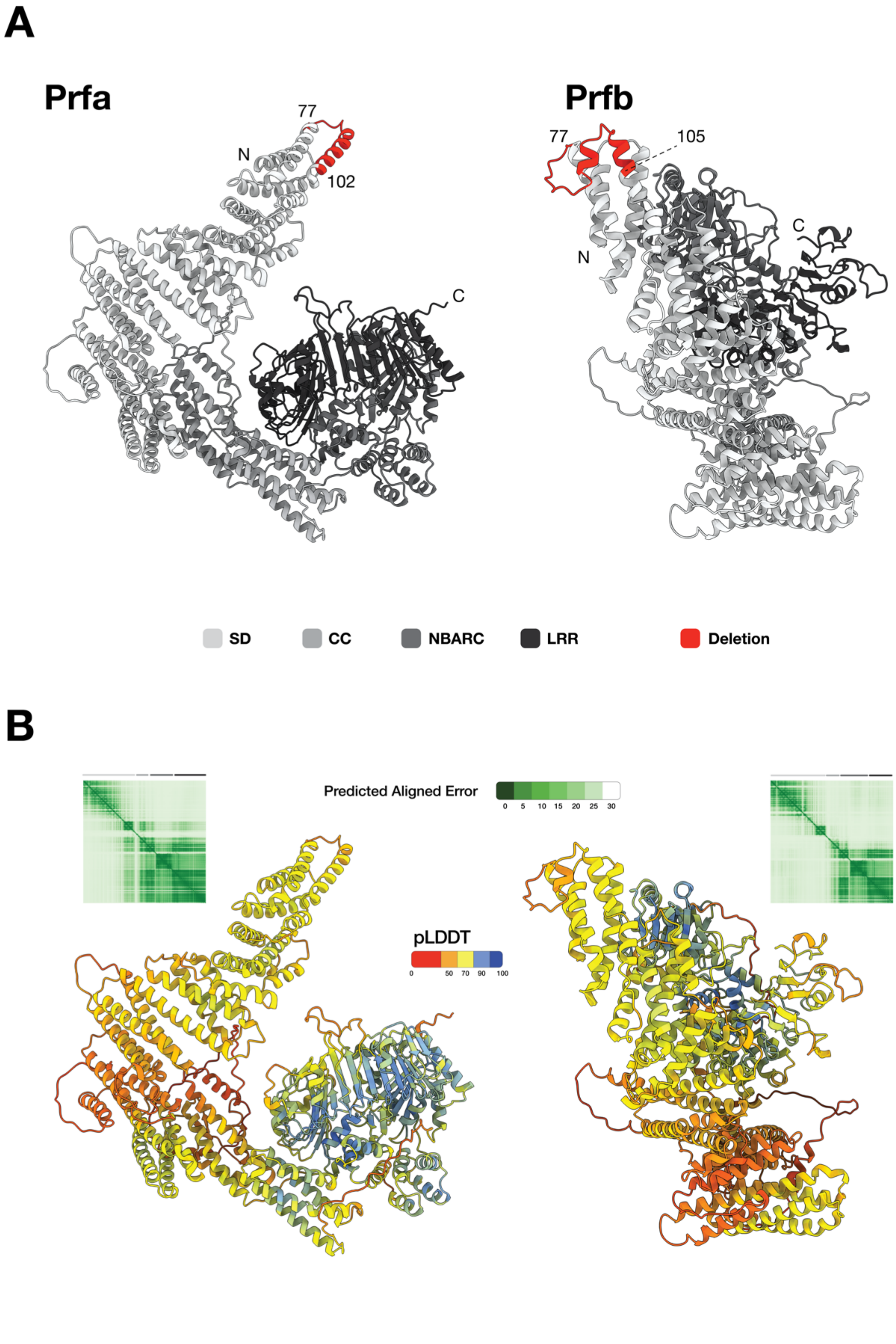


**Supplementary Figure 2: AlphaFold3 prediction of NbPrfa and NbPrfb.** The structures of Prfa and Prfb were predicted using AlphaFold 3 with a seed value of 1, based on the full-length amino acid sequences. Structure predictions were visualized using ChimeraX. **(A)** Different shades of gray are used to color the Solanaceous-domain (SD), Coiled-coil (CC) domain, nucleotide-binding adaptor shared by APAF-1, certain R gene products, and CED-4 (NBARC) domain, and the Leucine-rich repeat (LRR)

domain. The CRISPR/Cas9 target region within the CC domain is indicated in red. **(B)** The predicted Local Distance Difference Test (pLDDT) indicates the per-residue measure of local confidence, the Predicted Aligned Error (PAE) estimates the relative expected positional error for each residue. All data is available as Supplementary Dataset 5.

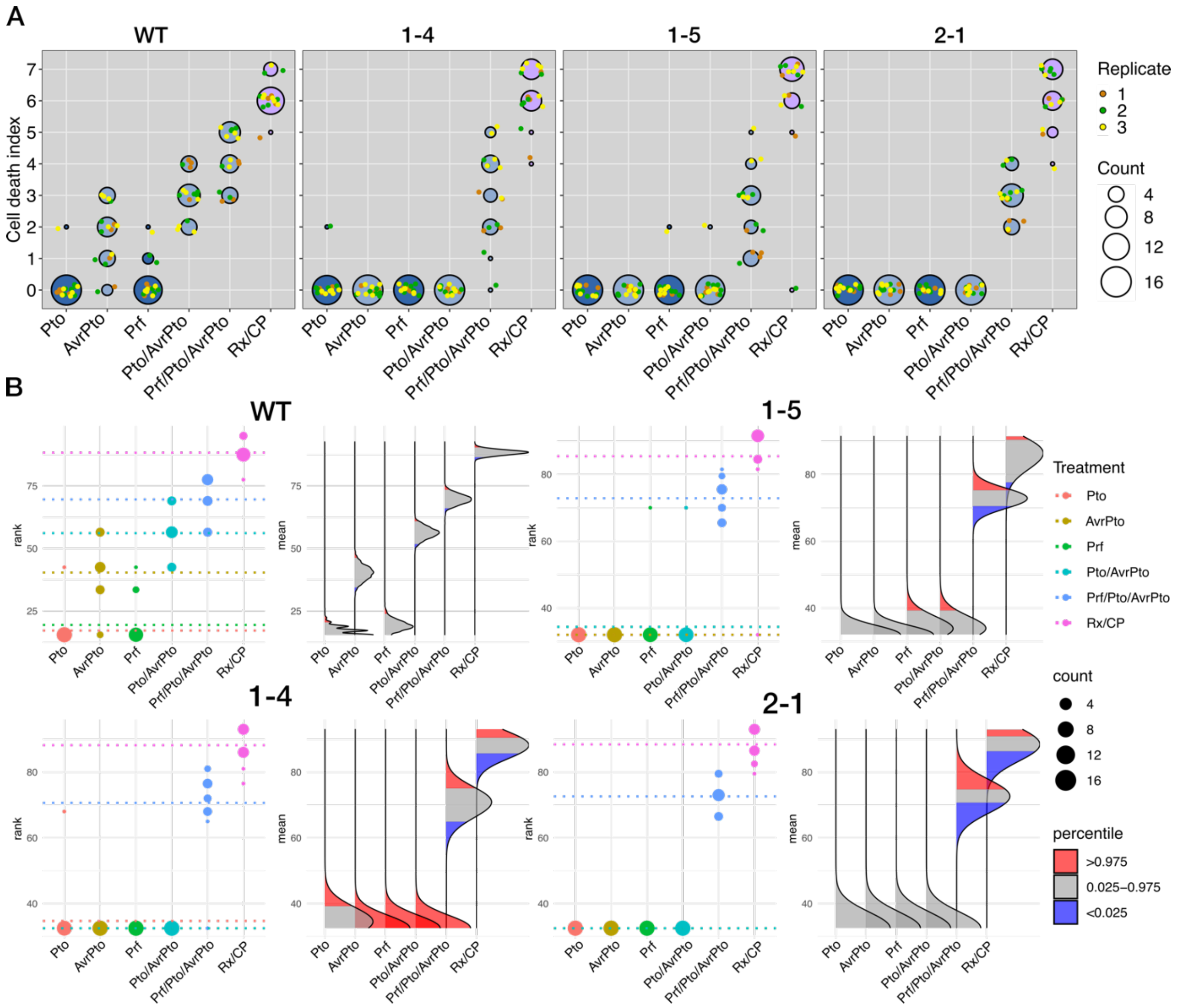


**Supplementary Figure 3: Cell death quantification and statistics for Figure 1B.** Cell death data is represented as dots, with different colors for biological replicates. The central circle for each cell death category proportionally indicates the total number of data points. Statistical analysis was performed using the besthr R library (MacLean, 2019). Data ranks are shown as dots with the corresponding mean as dashed line. The size of each dot proportionally indicates the total number of data points. A bootstrap resampling test was performed, using significance cutoffs of 0.025 and 0.975. Mean ranks of test samples falling outside of cutoffs in the control samples bootstrap population were considered significant. The distribution of 1,000 bootstrap sample rank mean is indicated, blue areas under the curve illustrate the 0.025, and red areas the 0.975 percentiles of the distribution. All data is available as Supplementary Dataset 4.

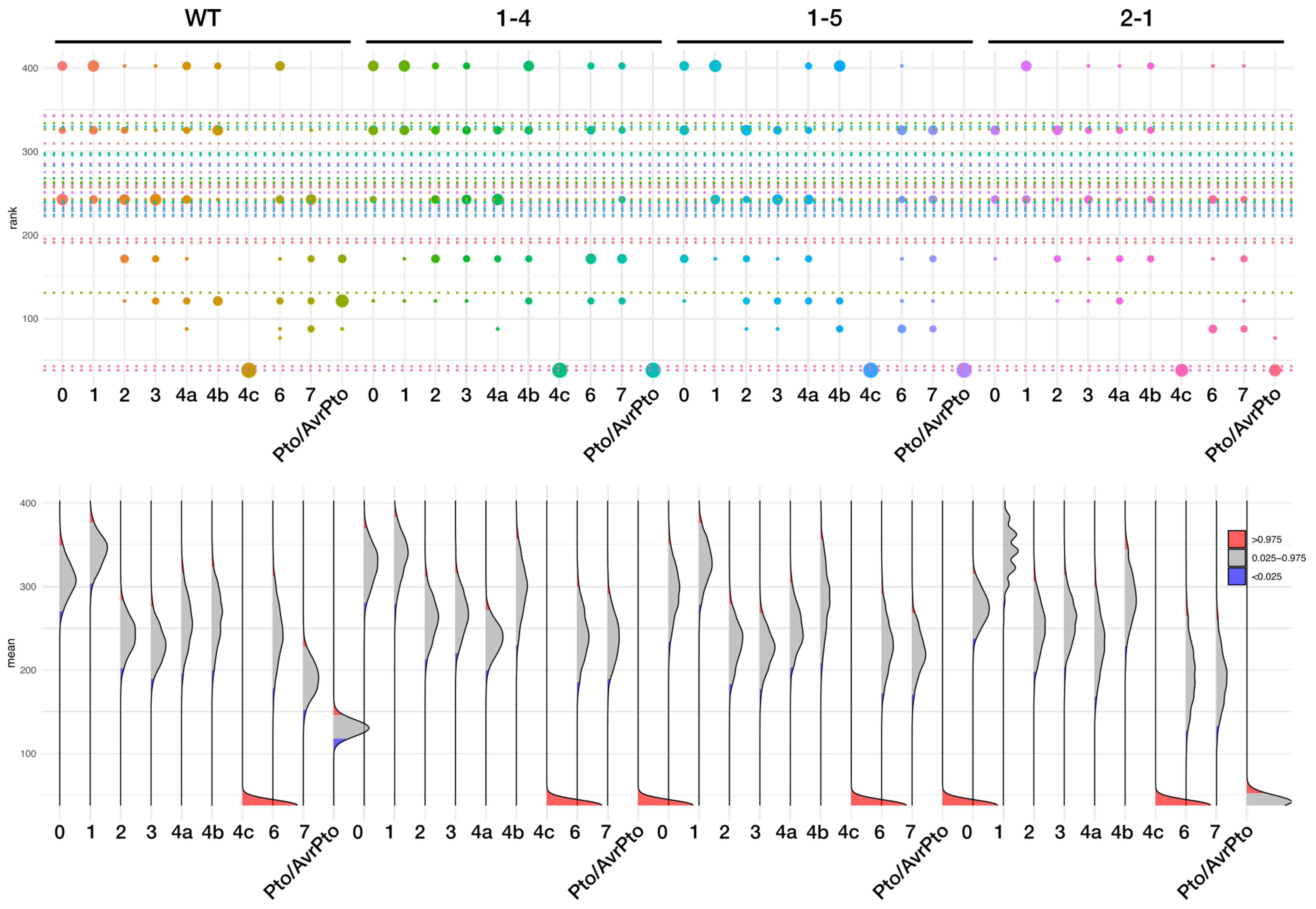

WT
1-4
1-5
2-1
rank
400
300
200
100
0 1 2 3 4a 4b 4c 6 7 Pto/AvrPto
mean
>0.975
0.025–0.975
<0.025

**Supplementary Figure 4: Cell death quantification and statistics for Figure 2.** Statistical analysis was performed using the besthr R library (MacLean, 2019). Data ranks are shown as dots with the corresponding mean as dashed line. The size of each dot proportionally indicates the total number of data points. A bootstrap resampling test was performed, using significance cutoffs of 0.025 and 0.975. Mean ranks of test samples falling outside of cutoffs in the control samples bootstrap population were considered significant. The distribution of 1,000 bootstrap sample rank mean is indicated, blue areas under the curve illustrate the 0.025, and red areas the 0.975 percentiles of the distribution. All data is available as Supplementary Dataset 4.
